# Multiblock LASSO Framework for Cancer Gene Selection from RNA-Seq PANCAN Data

**DOI:** 10.1101/2025.07.05.663323

**Authors:** Zeeshan Ashraf, Muhammad Aslam, Tahir Mehmood, Laila Abdulaziz Abdulrahman Al-Essa

**Author notes:** These authors contributed equally to this work.

## Abstract

The Cancer RNA-HiSeq PANCAN dataset consists of RNA-Seq gene expression data collected from multiple cancer types. It is a high-dimensional dataset, meaning it has thousands of gene expression features (predictors) and relatively fewer samples (observations). The dataset contains thousands of genes, making it difficult to identify key biomarkers. In order to reduce data and comprehend the modeled link, variable selection is essential. Least Regression using Absolute Shrinkage and Selection Operator (LASSO) is one modeling technique that deals with high throughput data. The data might be divided into different blocks representing different biological pathways or cancer types. Many genes are correlated, which can reduce interpretability. In many areas, including modern biology, variable selection is an important problem. For instance, choosing genetic characteristics for categorization (i.e., identifying harmful bacteria, diagnosing diseases, etc.) is an example of this. Multiblock Lasso (a variant of Lasso regression) is particularly useful when data is structured into blocks (e.g., different biological processes or pathways). It helps in selecting important features across multiple blocks, improving interpretability by grouping related genes, reducing over fitting in high-dimensional datasets. In this study, we apply Multiblock Lasso to extract significant gene features for cancer classification. We preprocess the dataset, define block structures using biological pathways, and optimize the regularization parameters using cross-validation. Experimental results demonstrate that Multiblock Lasso effectively reduces dimensionality while maintaining classification accuracy, making it a powerful tool for biomarker discovery in cancer genomics.

## 1. Introduction

This study draws on a curated portion of the RNA-Seq HiSeq Pan-Cancer dataset. There are featuring gene-expression profiles from 801 tumor samples representing five distinct cancer types, Lung adenocarcinoma (LUAD), Breast invasive carcinoma (BRCA), Colon adenocarcinoma (COAD), Prostate adenocarcinoma (PRAD) and Kidney renal clear cell carcinoma (KIRC). The dataset comprises 20,531 gene features, capturing high-dimensional transcriptomic variations, and is suited for multivariate analysis, particularly classification and clustering tasks in cancer genomics. Researchers increasingly favour single-cell RNA sequencing for measuring gene activity in cancer and other diseases because it produces cleaner, less-studied noise and can discover new transcripts without needing a preset list of abundant genes [1]. In large single-cell RNA-seq studies, speedy and reliable feature selection is key to good classification. Pulling out the most informative genes from thousands of candidates sharpens explanation, boosts prediction, and cuts the time and money needed to take scRNA-seq into clinics, drug discovery, and genetic screens [2]. An imbalanced classification problem pops up whenever some classes in the dataset have far more examples than others. Working with small, uneven training sets can be especially tough in fields like bioinformatics or medical research, where every sample can be hard to come by yet carries a lot of meaning. If these situations are not handled carefully, the resulting model may treat rare classes unfairly, focus too much on obvious features, and end up missing the real, subtle signals that tell the story hidden in the data [3]. Analysis using ribonucleic acid sequencing (RNA-Seq) is very helpful for learning more about genes that are expressed differently. Its high-dimensional data, however, makes it difficult. This type of analysis is a method for identifying underlying patterns in data, such as biomarkers unique to cancer. RNA-Seq data related to the same cancer class as positive and negative samples that is, without samples of other cancer types were analyzed in the past. [4]. Cancer is a highly complex and heterogeneous disease. A machine learning workflow consisting of dataset identification, normalization, feature se- lection, dimensionality reduction, clustering, and classification was implemented [5]. Explainable Machine Learning (XML) approaches are crucial for medical information processing tasks, particularly for multi-omics data analytics. The XML system not only provides better performance but also explains the inside of the finding better. Here, we proposed an end-to-end explainable system for analyzing high dimensional RNA-seq data using an unsupervised gene selection approach and supervised methods, including Deep Neural Network (DNN) [6]. Feature selection criteria / techniques such as in gene marker identification and cell type categorization. The most widely used techniques for selecting features from scRNA-seq data are predicated on the idea of differential distribution, which uses a statistical model to identify variations in gene expression between cell types [7]. This work uses omics data analysis and computational techniques to address the urgent need for better lung cancer detection and therapy. More efficient diagnosis and treatment methods are urgently needed since lung cancer continues to be a major cause of cancer-related deaths worldwide [8].This is significant from a modeling perspective, since it improves the precision of identifying the reality underlying a behavior by integrating biological data on genes with comparable functions [9]. Using the average of the variables to represent a group of associated variables is one method of selecting a set of variables. The variable selection model can then only employ this representative member. When averaging variables, bias in the variable selection process might exist. Ideal subset selection produces an unbiased model, but lasso usually selects individual variables instead of groups [10, 11]. It is recommended to use LASSO regression to determine the important variables within each group, and then group lasso to choose relevant clusters. Elastic net selects entire clusters of variables by including all related variables when one is chosen [12]. However, it is only possible to choose a cluster of strongly interrelated variables if their regression coefficients are nearly equal. For cluster selection, this encourages the use of a strong structure extraction technique. LASSO is widely used for high-dimensional data to manage cases with more variables than samples and multicollinearity [13]. Numerous advancements in LASSO have been developed to accommodate various data structures. One such method, Multiblock LASSO (mbLASSO), is particularly suited for scenarios where groups of variables can be organized into blocks or clusters. When it comes to selecting relevant blocks, two main approaches are commonly used. One approach involves conducting statistical significance tests on the coefficients within each block [14] and the other approach relies on evaluating the importance of each block in terms of its contribution to the predictive performance, commonly referred to as Block Importance on Prediction (BIP) [15]. A recent study suggests selecting clusters based on their stability in order to achieve more reliable and interpretable results. [16]. Building on this idea, we employed a modified version of the regularized stepwise procedure [17], which allows for the elimination of a substantial number of clusters with only a negligible increase in the model’s root mean square error (RMSE). This trade-off enhances both the interpretability of the model and the stability of the variable selection. In this approach, clusters are ranked using the Block Importance on Prediction (BIP) measure within the stepwise framework. We demonstrate the effectiveness of this procedure through an application focused on identifying codon and di-codon usage patterns associated with optimal growth temperatures in prokaryotes.

## 2. Approach

### 2.1 Data

The data collection is a random extraction of gene expressions from patients with various tumor types, including BRCA, KIRC, COAD, LUAD, and PRAD (*https://archive.ics.uci.edu/dataset/401/gene+expression+cancer+rna+seq*). It is a component of the RNA-Seq (HiSeq) PANCAN data set where samples (instances) are stored row-wise. Each sample’s RNA-Seq gene expression levels, as determined by the Illumina HiSeq platform, are its variables (attributes). Numerous attempts have been made to use gene expression data to improve the accuracy of cancer categorization. However, using large amounts of data raises the risk of data overfitting, which emphasizes the need for more effective methods. In order to make the dataset usable, we have employed mild forms of dimensionality reduction to lower the amount of features. The RNA-Seq (HiSeq) PANCAN dataset from the UCI Machine Learning Repository, which includes a comprehensive collection of gene expression data across several tumor samples, is used in this work to investigate the classification of different cancer types. This study uses a dataset consisting of randomly selected gene expression profiles from patients diagnosed with different tumor types. This data, sourced from The Cancer Genome Atlas (TCGA) and hosted by the UCI Machine Learning Repository, contains high-throughput RNA-Seq measurements that capture transcriptomic variations across diverse cancer types. These profiles enable comprehensive analyses for identifying discriminative gene patterns, facilitating biomarker discovery, and enhancing classification or prediction models in oncology-related research.

### 2.2 Clusters of Variable

Before fitting any models, by deducting the column mean and dividing by the standard deviation, all variables in both y and x were centered and standardized. To assess similarity between variables, the correlation-based pairwise distance d was calculated for each pair of variables x_?_ and x_j_ as,

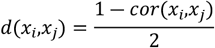

Here, cor represents the correlation between x_?_ & x_j_, and the resulting distance d is a value between 0 and 1. Subsequently, every variable was shown as a node in an undirected network. An edge was drawn between two nodes if the distance between them was less than a specified threshold t. A smaller t ensures that only highly correlated variables are connected. This graph structure naturally forms C clusters, with each cluster X^c^ (for c = 1, 2, …, C) containing P_c_ variables. This defines *X* = [*X*^*1*^*…*..*X*^*C*^] having *p* = Σ *p*_*c*_ columns.

### 2.3 Multiblock Lasso

This is an adaptation of the traditional Lasso regression designed to handle data organized in blocks, where each block represents a group of related variables. This approach applies a Lasso penalty separately to each block, enabling simultaneous variable selection and estimation within each block while taking into account the overall multiblock structure. The relationship between the response variable y and the different blocks *X* = [*X*^(1)^,*X*^(2)^…. *X*^(*C*)^] is assumed to be linear. Since we have to approach Lasso penalty separately to each block, this can be addressed using mbLASSO. The primary goal of this algorithm is to find scores for each block, denoted as s, which are then used to create combined scores t that illustrate the relationship between blocks.. Here, we have adopted the mbLasso procedure with some modification, where loading

Algorithm starts with *E*_0_ = *X* = [*X*^(1)^,*X*x^(2)^…. *X*^(*C*)^] and *f*_0_ = 𝒴.

For *r* = 1 *to R*

for *c* = 1 *to C*

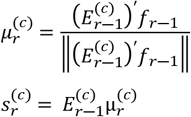

*l*(*c*) = *number of columns of E*^(*c*)^

*end*

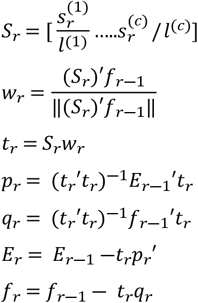

*Extract each block* 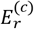*from E*_*r*_

*end*

For prediction, model coefficients are stored; 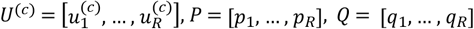 and *W* = [*w*_1_, …, *w*_*R*_], and for test data *N* = [*N*^(1)^, …, *N*^(*C*)^ which is scaled as X with ŷ = 0 and *E*_0_ = *N*.*c*

For *r* = 1 *to R*

for *c* = 1 *to C*

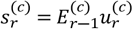

end

*Extract each block* 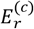*from E*_*r*_

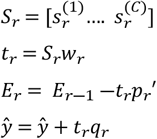

end

### 2.4 Training-Test Splits and Nested Parameter Tuning in mbLASSO

In this study, a three-level model validation and tuning strategy was employed to enhance the accuracy and generalizability of the multiblock LASSO (mbLASSO) approach for gene expression data analysis. At the first level, the dataset was randomly split 30 times into 75% training and 25% test sets, with partial overlap, and the RMSEP (Root Mean Square Error of Prediction) was calculated for each split to assess predictive performance. Within each training set, a second level of 10-fold cross-validation was used to optimize two key parameters: the step length *b ∈* {0.1,0.5,1}, and the rejection threshold *m* ∈{0.90,0.99}, while fixing the regularization parameter *α* =10. In the context of your study, LOOCV was used in the last stage to fine-tune and estimate the best parameters for the model by repeatedly testing its accuracy on every individual data point. This helps ensure that the model is robust and generalizes well to new data. This nested and repeated validation structure forms a robust cross-model validation framework, ensuring both stable feature selection and reliable prediction performance.

### 2.5 Algorithm for cluster selection

We recently introduced a stepwise estimation method aimed at selecting a minimal set of relevant variables. The data has been randomly divided into a predetermined number of subsets (test and training) using a stability-based variable selection technique. The variables are chosen in a stepwise manner for every split. Ultimately, stable variables are chosen by a process of stepwise reduction from every data split. This approach was also used here, although feature selection was done on groups of variables rather than on individual variables, and a greedy algorithm was used to repeatedly remove the “worst” clusters. The blocks in X must be ranked in order for the algorithm to work. Block importance on prediction (BIP) is used for this and is defined as

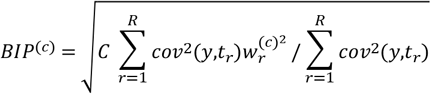

The contribution of each block to these components via weights are 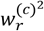. The correlation between latent components *t* _*r*_ and the response 𝒴. BIP quantifies the relative predictive importance of each block X^(c)^. *W* represents loading weights of the cluster c and cov means covariance. Each cluster’s contribution is weighted by the BIP based on the variation that each PLS component explains. Cluster c can be eliminated, if *BIP*^(*c*)^< *α* for some user-defined threshold *α* ∈ [0,∞). Defining *α* is a critical issue, here we have modified the stepwise algorithm [18] for cluster selection.

The stepwise cluster elimination algorithm can be sketched as follows: Let *Z*_0_ = *X* = [*X*_1_,*X*_2_,……,*X*_*C*_].

1. For iteration *g* run *y* and *Z*_*g*_ through cross validated mbLASSO. The matrix *Z*_*g*_ has *p*_*g*_ clusters, and we get the same number of criterion values, sorted in ascending order as BIP^(1)^,….BIP^(Cg)^.
2. These are M criterion values below the cutoff *α*. If *M=0*, terminate the elimination here.
3. Else, Let *N* = [*bM*] for some fraction *b* ∈ < 0,1]. Eliminate the clusters corresponding to the N most extreme criterion values.
4. If there are still more than one cluster left, Let *Z*_*g+1*_ contain these clusters, and return to 1. The “steplength” of the elimination procedure is determined by the fraction b; a b near 0 will only remove a small number of clusters in each iteration. Figure 2 provides an overview of block removal. Cross validation can be used to determine the fraction *b* and threshold *a*.

**Figure 1:**
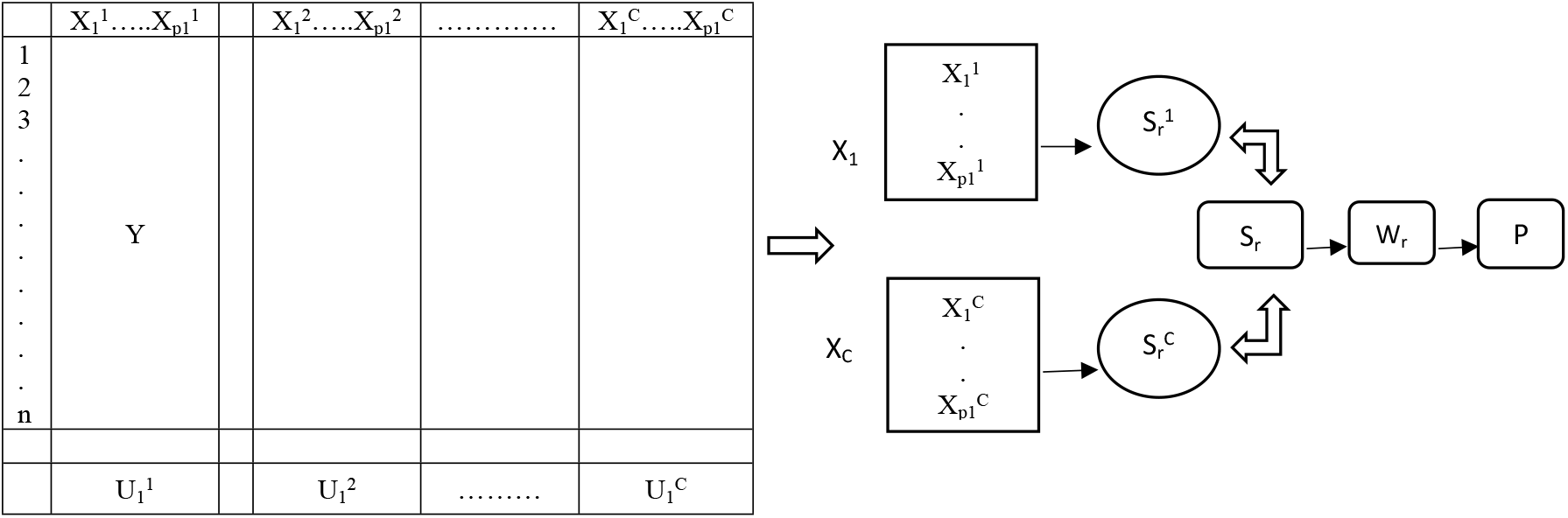
A summary of the relationship between clusters and latent variables (mbLASSO).

**Figure 2.**
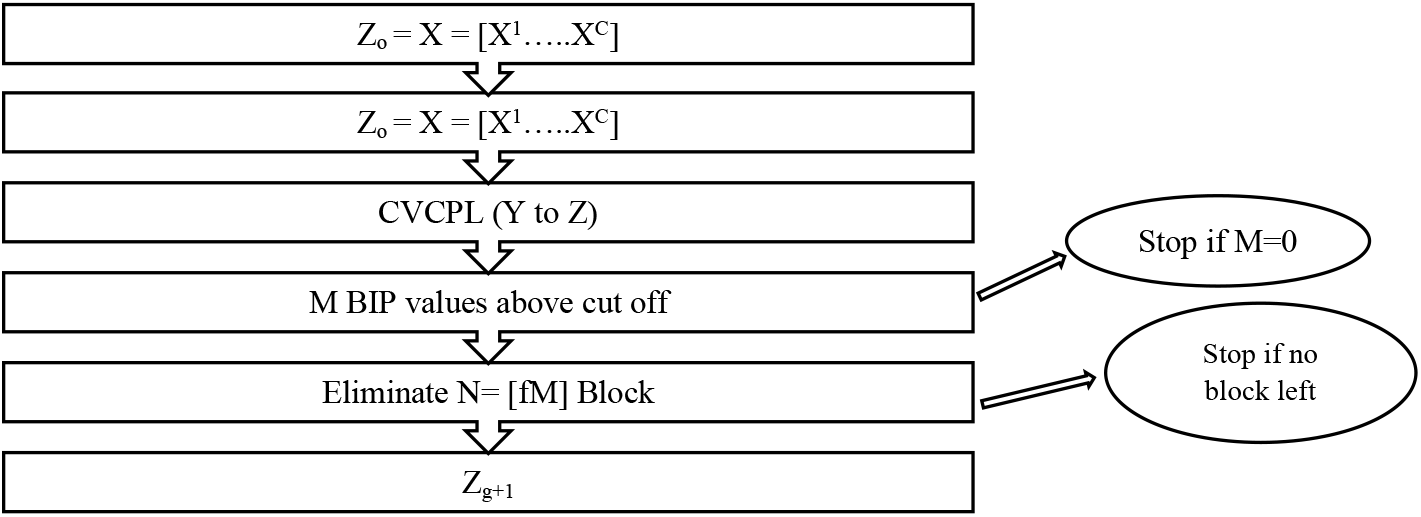
A summary of the block eliminatiom strategy employed in our stepwise elimination procedure.

**Figure 3.**
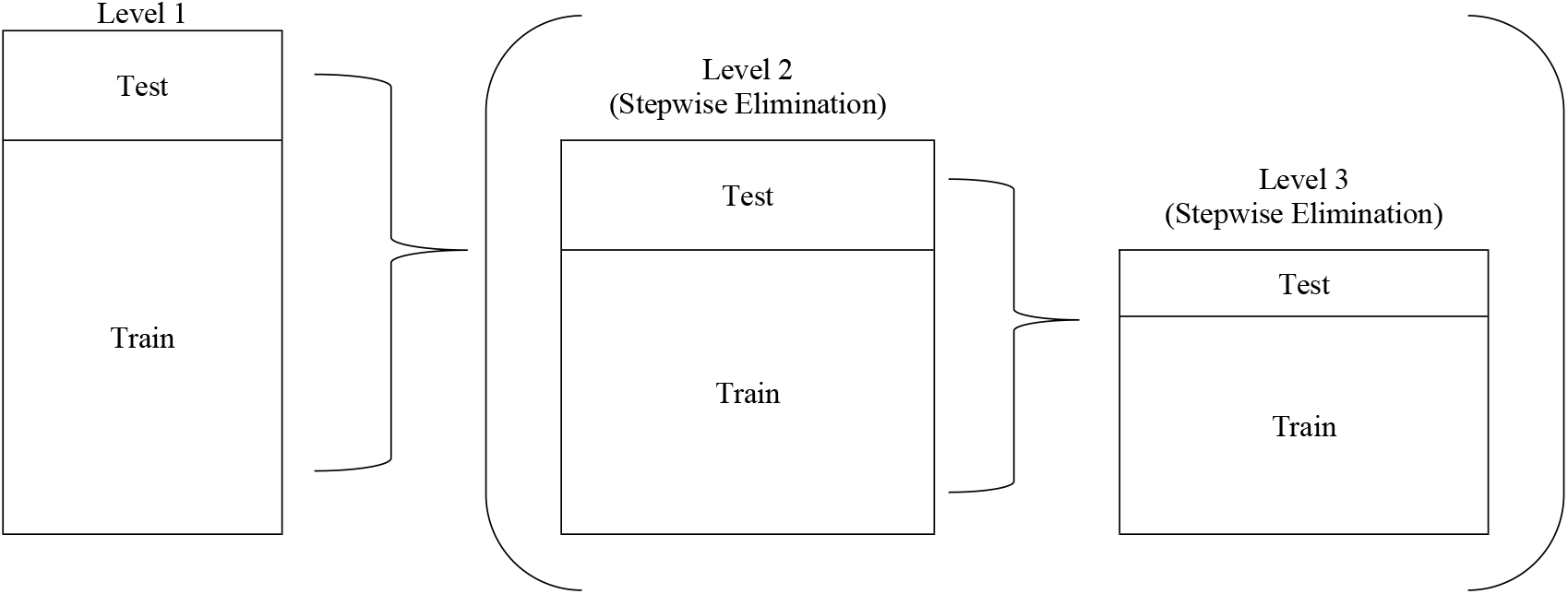
An outline of the training-testing process employed in this investigation is provided. The predictor matrix is represented by the rectangles. To guarantee robustness, the data were randomly divided into a training set (75%) and a test set (25%), and this procedure was carried out 30 times at Level 1. We added two more stages of cross-validation to our suggested stepwise reduction strategy. The selection parameters “a” and “b” were optimized at Level 2 using a 10-fold cross-validation. At Level 3, the regularized mbLASSO model was refined using leave-one-out cross-validation.

We obtain the root mean square error (RMSE) for every iteration g of the elimination, which is represented by *L*_*g*_. The prediction ability shows good represents by a RMSE value near zero. With every iteration, the number of impacting clusters diminishes, also *L*_*g*_ decreases until nearly optimum is achieved. The values will increase again as we keep on eliminating. For a relatively minor compromise of optimal RMSE, a potentially much simpler model may be obtained. Accordingly, we require a rejection threshold *m*, where each iteration that goes over the ideal RMSE *L*^*\**^. To provide insight into the trade-off between the model’s understandability and RMSE, we may calculate the t-test and p-value.

## 3. Results and Discussion

We have used the mbLASSO to classify DNA samples based on various cancer types using data from the UCI Repository. The dataset titled “Gene Expression Cancer RNA-Seq” from the UCI Machine Learning Repository has the following dimensions: Samples (Patients): 801, Features (Genes): 20,531 gene expression values per sample. This dataset is unbalanced, as the number of samples varies significantly across cancer types for example, breast cancer has the highest count with 300 samples, while colon cancer has the lowest with only 78. Despite this imbalance, multiple gene groups were identified that enable classification with reasonable accuracy. These gene groups function as individual classifiers, and their outputs are integrated using a voting-based decision system. All numerical results reported from the experiments represent averages over 500 independent and randomly initialized iterations.

Table 1 summarizes the distribution of samples across five different tumor types in the dataset. The largest group consists of 300 samples from Breast Invasive Carcinoma (BRCA), followed by Kidney Renal Clear Cell Carcinoma (KIRC) with 146 samples, Lung Adenocarcinoma (LUAD) with 141 samples, Prostate Adenocarcinoma (PRAD) with 136 samples, and Colon Adenocarcinoma (COAD) with 78 samples. In total, the dataset includes 801 samples, representing a diverse set of cancer types for analysis. The Lasso model is initialized with a specified regularization strength, denoted by the parameter λ. This parameter plays a crucial role in controlling the complexity of the model by adding a penalty term to the loss function, which discourages excessively large coefficients.

**Table 1.** Division of DNA microarray data of UCI Repository.

| Number of samples | Class ID | Class |
| --- | --- | --- |
| 300 | 1 | BRCA |
| 136 | 2 | PRAD |
| 146 | 3 | KIRC |
| 141 | 4 | LUAD |
| 78 | 5 | COAD |
| 801 |  |  |

**Table 2:** The values generating from 10 iterations of different lambda values by the procedure of mbLASSO.

| $\lambda$ | Opt $\lambda$ | Selected Variables |
| --- | --- | --- |
| 0.1 | 0.098 | 386 |
| 0.2 | 0.076 | 321 |
| 0.3 | 0.0306 | 303 |
| 0.4 | 0.022 | 292 |
| 0.5 | 0.0183 | 278 |
| 0.6 | 0.0152 | 263 |
| 0.7 | 0.031 | 261 |
| 0.8 | 0.011 | 258 |
| 0.9 | 0.0102 | 254 |

This plot represents the LASSO (Least Absolute Shrinkage and Selection Operator) regression path in Fig. (4), showing how coefficients of different variables change as the regularization parameter (lambda) varies. The model is then fitted to the training data, enabling it to learn the underlying patterns present in the dataset. Dimensionality of features is an important factor for any multivariate analysis. Selecting clusters based on their stability can lead to more meaningful and reliable interpretations. To assess model stability and selectivity, we recently proposed a straightforward selectivity score. when a variable is chosen among *p* variables, it receives a score of *1/p*. X-axis (Log Lambda 0.1 -0.9), represents the log-transformed value of the regularization parameter (λ). Moving left (lower λ): Less regularization, more variables included. Moving right (higher λ): Stronger penalty, more coefficients shrink to zero.

**Figure 4.**
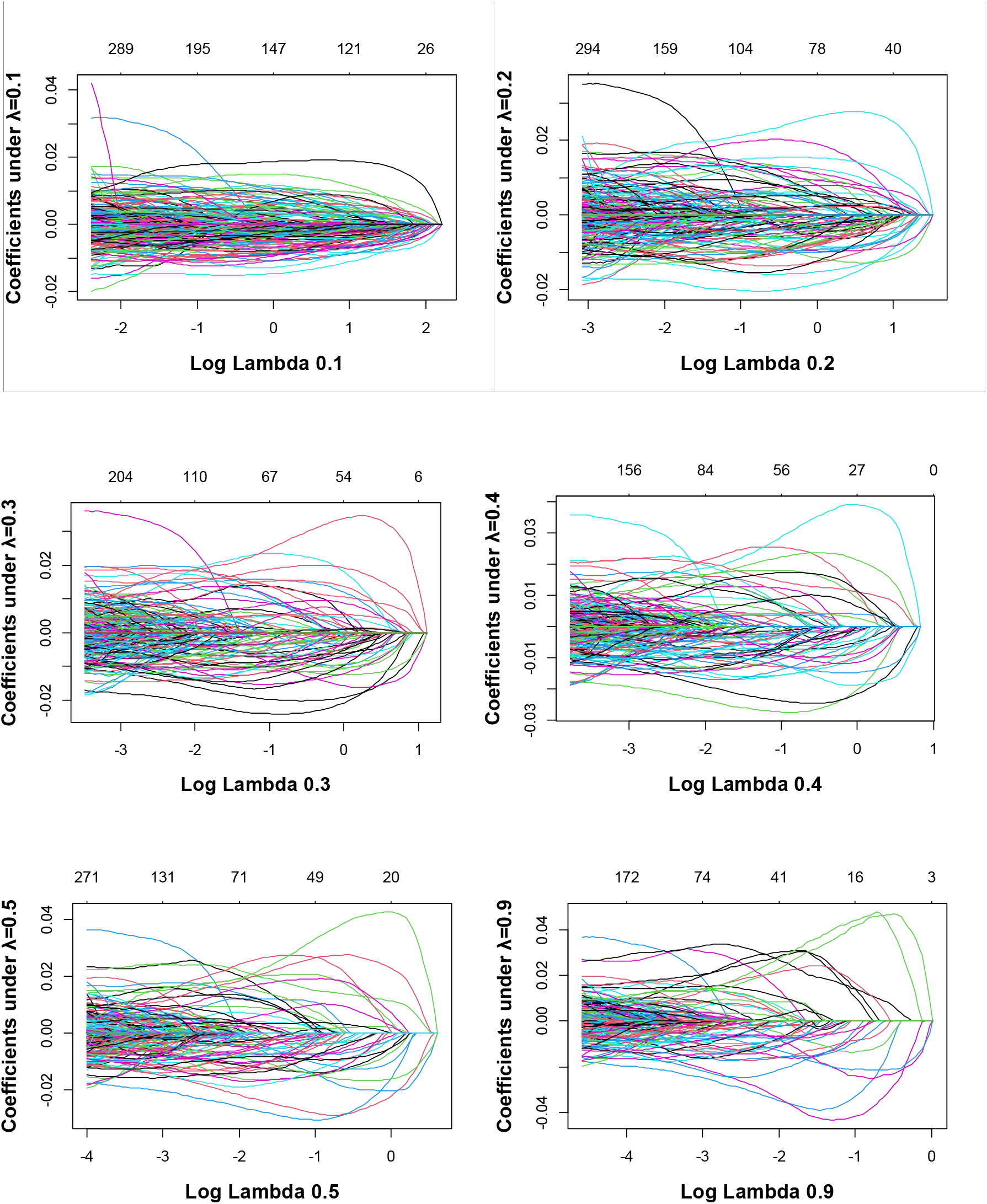
Coefficients of variable in accordance λ=0.1, 0.2, 0.3, 0.4, 0.5, 0.9. Complete set of coefficients according to the concerned lambda value

Y-axis (Coefficients under λ = 0.2), displays the values of the regression coefficients for different features. Many lines start large and shrink toward zero as λ increases. Few features remain nonzero at high λ, indicating strong predictors. Variables with coefficients close to zero early on may not be significant for prediction. Next we have to choose an optimal λ (based on cross-validation or a specific threshold). Features with nonzero coefficients at optimal λ are selected (important predictors). Eliminate features with coefficients shrinking to zero early on to reduce model complexity.

In this plot of Mean-Squared Error (MSE) versus Log(λ), likely from a cross-validation process in a mb-LASSO. The x-axis represents the Shrinkage parameter log (Log(λ)). The y-axis represents the Mean-Squared Error (MSE). The red points indicate MSE values for different values of λ. The error bars represent the variability (standard error) in MSE. The top axis values likely correspond to the number of nonzero coefficients in the model at each λ. The two vertical dotted lines might indicate the optimal λ values, commonly the minimum MSE (left) and the one-standard-error rule selection (right). λ_min (Left Line): Produces the most accurate model with minimal MSE. Retains more features (less regularization). λ_1se (Right Line): Provides a simpler model with fewer features but still within acceptable error limits. Often preferred for better generalization (reduces over fitting).

The Classification for variables under lambda 0.1 is three hundred sixty eight variables. These variables are effective for the RNA-Seq gene expressions. When we have lambda in minimum position then it will be shown that highest no of features are directly effects positively or negatively to the gene expression. This plot suggests that only a few variables contribute significantly, and many features could be discarded without losing much information. This is useful for feature selection, reducing dimensionality, and improving model efficiency. This scatter plot likely represents the importance scores of features after applying a variable selection method (e.g., LASSO, Ridge, or another feature selection technique). X-axis (Feature Index): Each point represents a different feature from the dataset, indexed numerically. Y-axis (s0 - Feature Importance Score or Coefficient Value): Indicates the importance or contribution of each feature to the model. Values close to zero suggest the feature is less relevant. Higher values indicate stronger influence in the model.

The feature selection in our study, so as the value of lamba increases the optimal lambda values gets decreasing as shown in Figure 5. Higher values of lambda apply a stronger penalty, causing more coefficients to shrink toward zero and thereby decreasing the influence—or completely removing—certain features from the model, which enables automatic feature selection. In contrast, lower lambda values lessen the penalty’s impact, allowing more features to be retained in the model.

**Figure 5.**
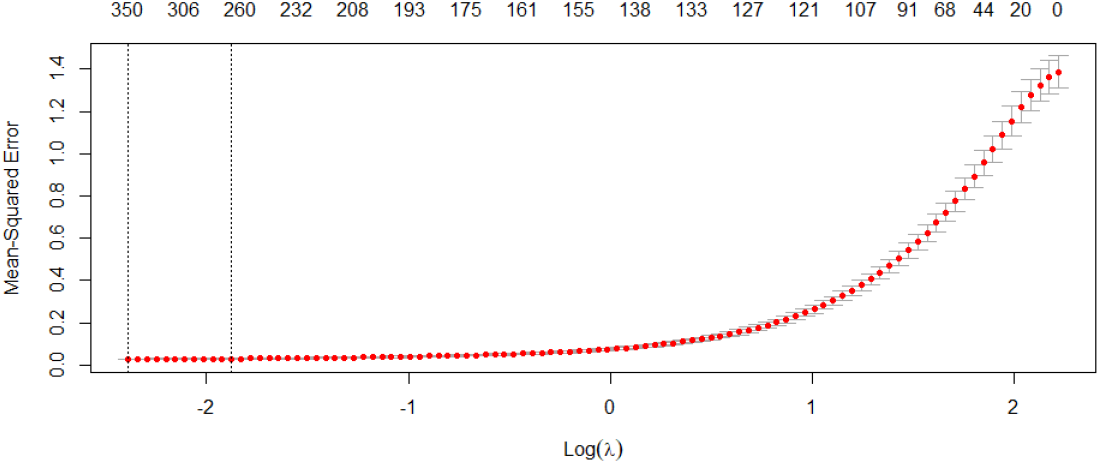
The mean squared error using K-fold cross-validation to suggest a lambda value of 0.1 for the lasso model.

**Figure 6.**
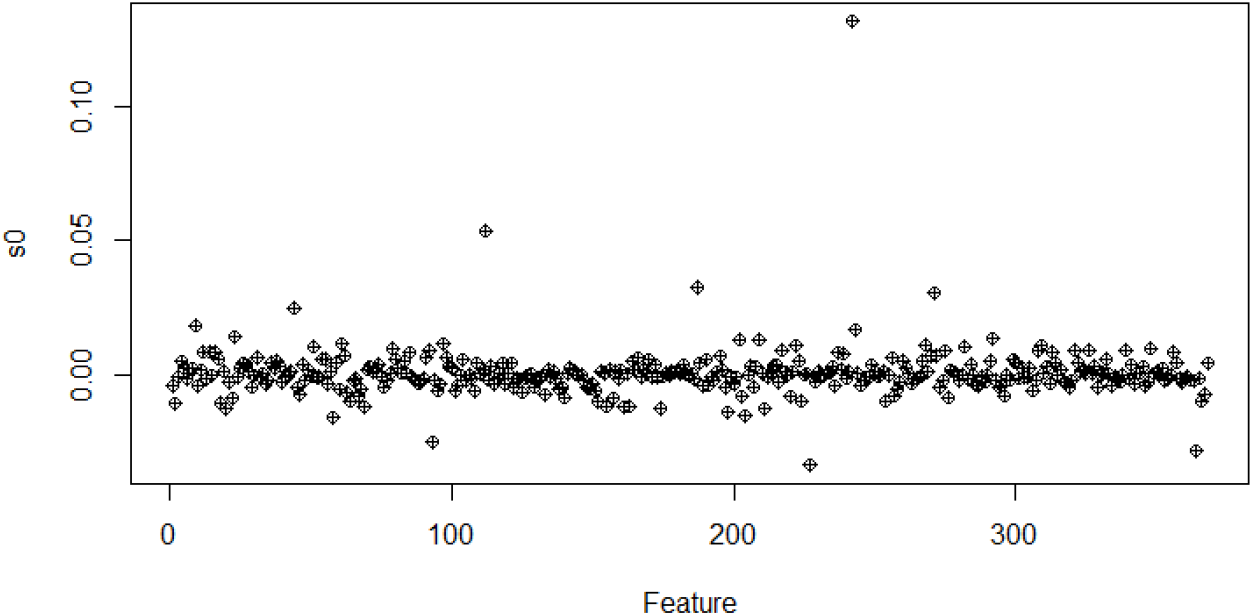
Classification of the variables by mblasso regarding Lambda = 0.1 is 368

**Figure 7.**
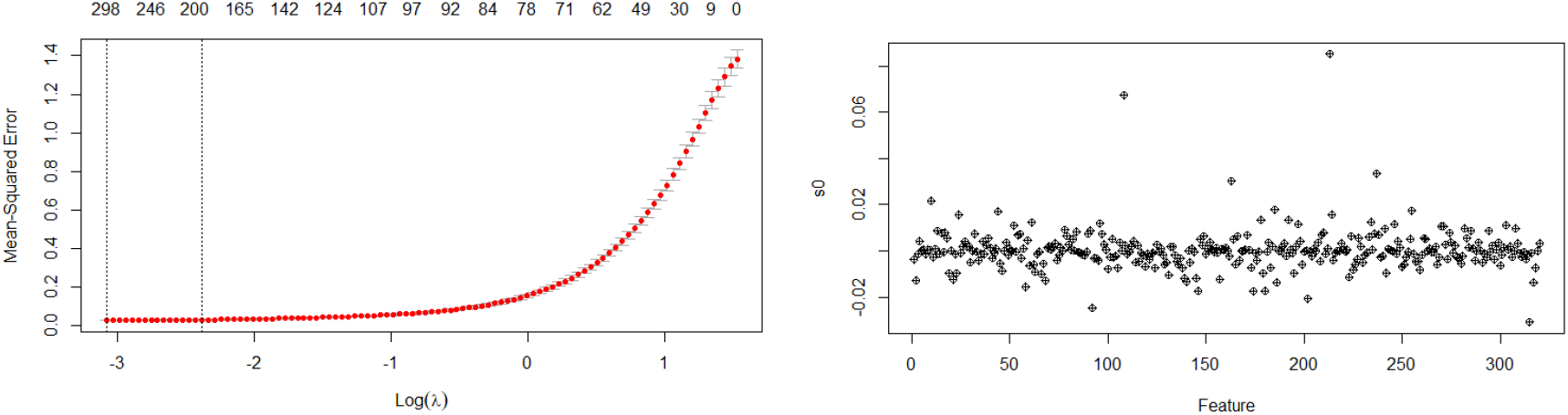
Calculates the mean squared error using K-fold cross-validation to suggest a lambda value for LASSO. Classification of the variables by mbLASSO regarding Lambda = 0.2 is 321.

**Figure 8.**
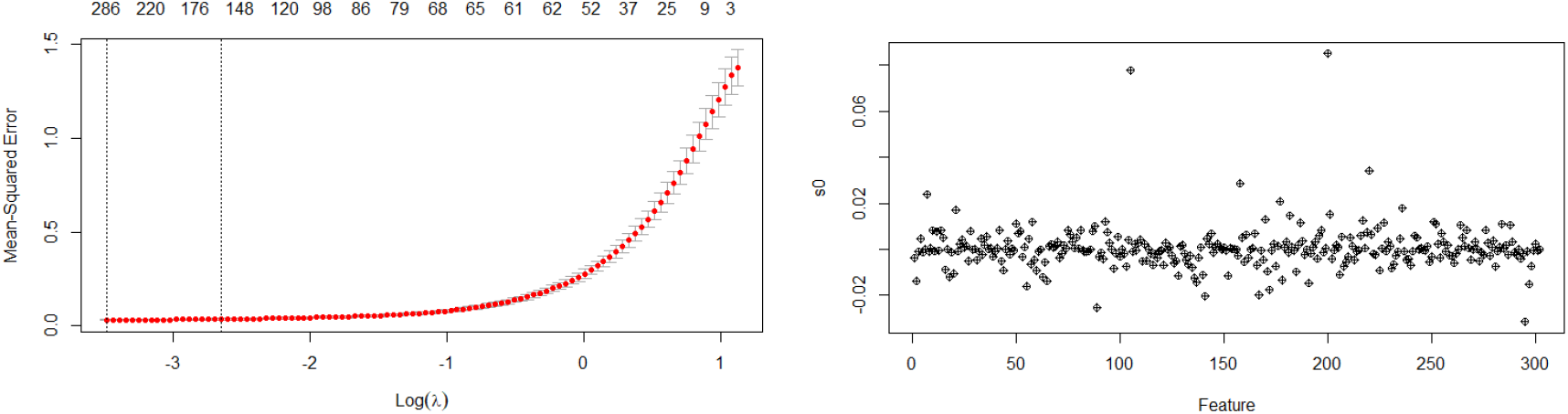
Calculates the mean squared error using K-fold cross-validation to suggest a lambda value for LASSO. Classification of the variables by mbLASSO regarding Lambda = 0.3 is 303.

**Figure 9.**
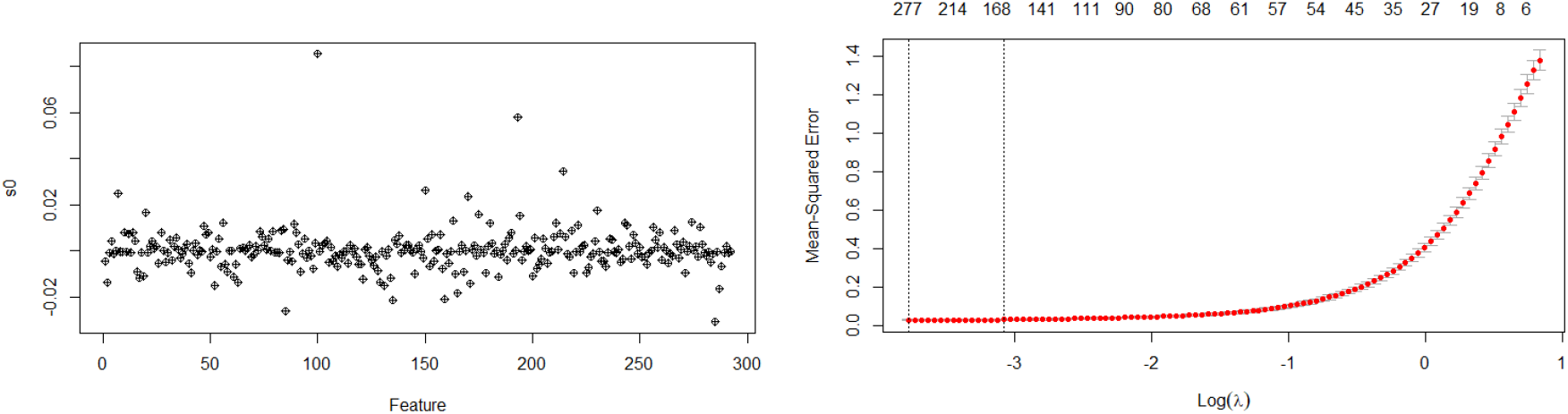
Calculates the mean squared error using K-fold cross-validation to suggest a lambda value for LASSO. Classification of the variables by mbLASSO regarding Lambda = 0.4 is 292.

**Figure 10.**
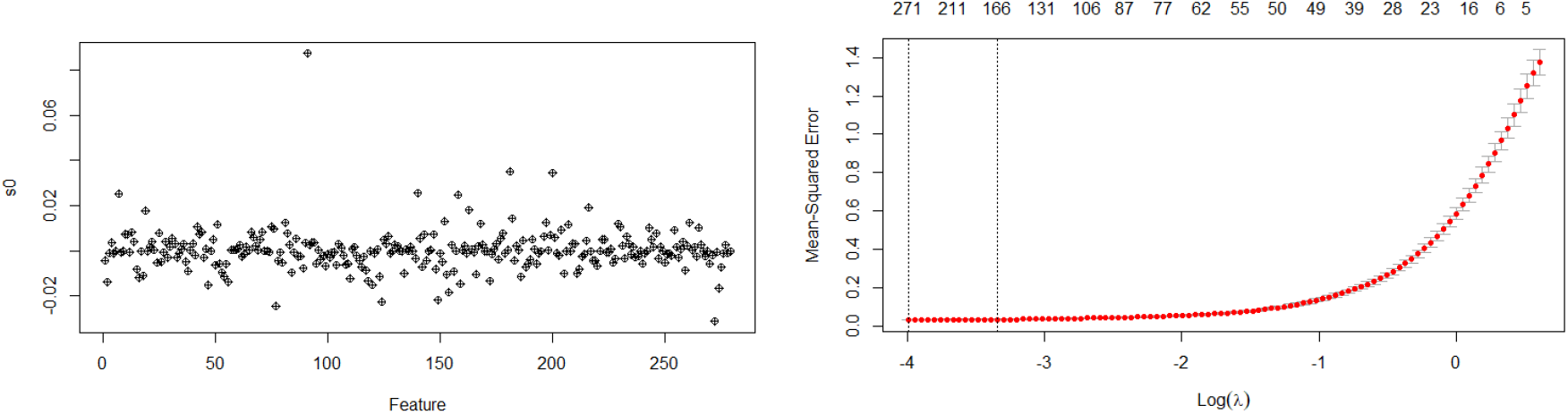
Calculates the mean squared error using K-fold cross-validation to suggest a lambda value for LASSO. Classification of the variables by mbLASSO regarding Lambda = 0.5 is 278.

**Figure 11.**
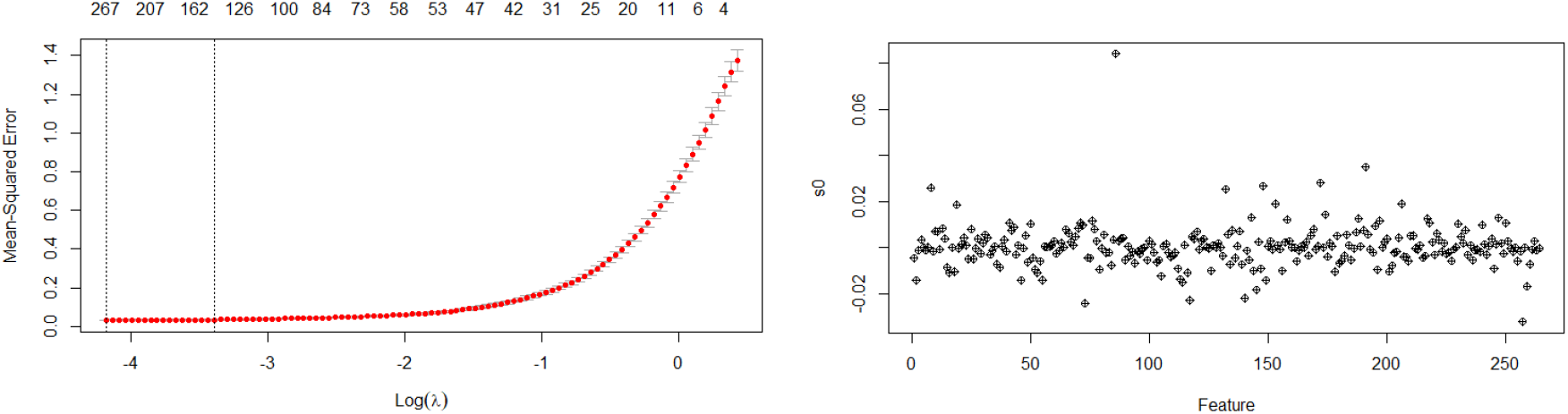
Calculates the mean squared error using K-fold cross-validation to suggest a lambda value for LASSO. Classification of the variables by mbLASSO regarding Lambda = 0.6 is 263.

**Figure 12.**
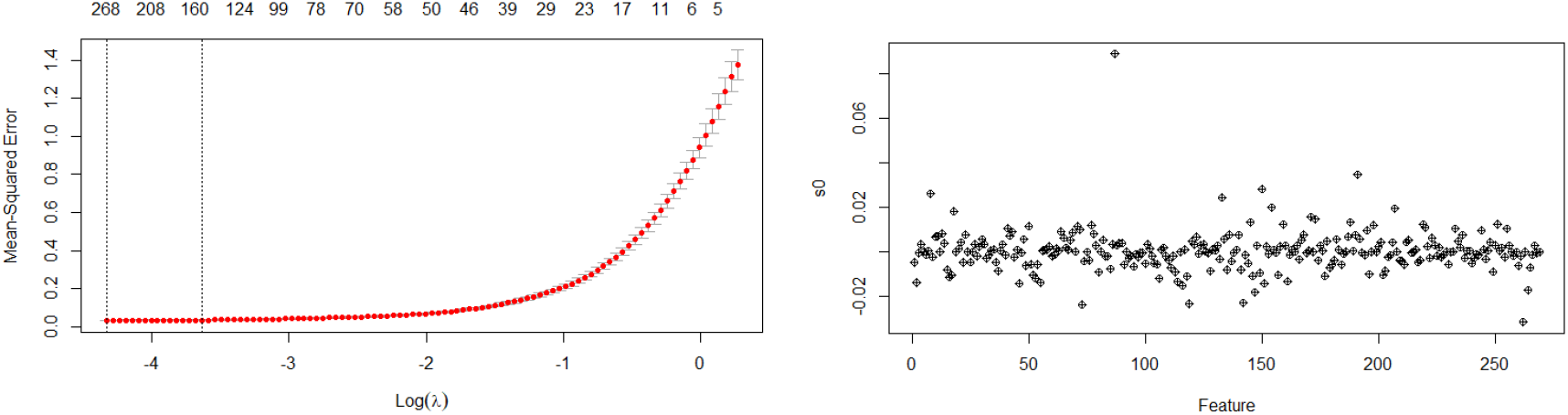
Calculates the mean squared error using K-fold cross-validation to suggest a lambda value for LASSO. Classification of the variables by mbLASSO regarding Lambda = 0.7 is 261.

**Figure 13.**
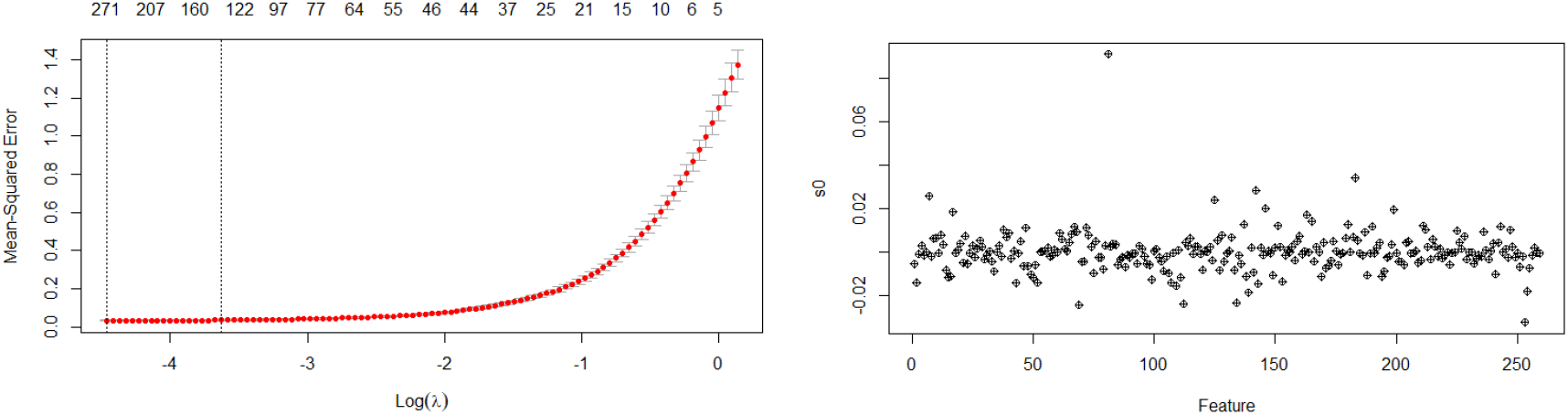
Calculates the mean squared error using K-fold cross-validation to suggest a lambda value for LASSO. Classification of the variables by mbLASSO regarding Lambda = 0.8 is 258.

**Figure 14.**
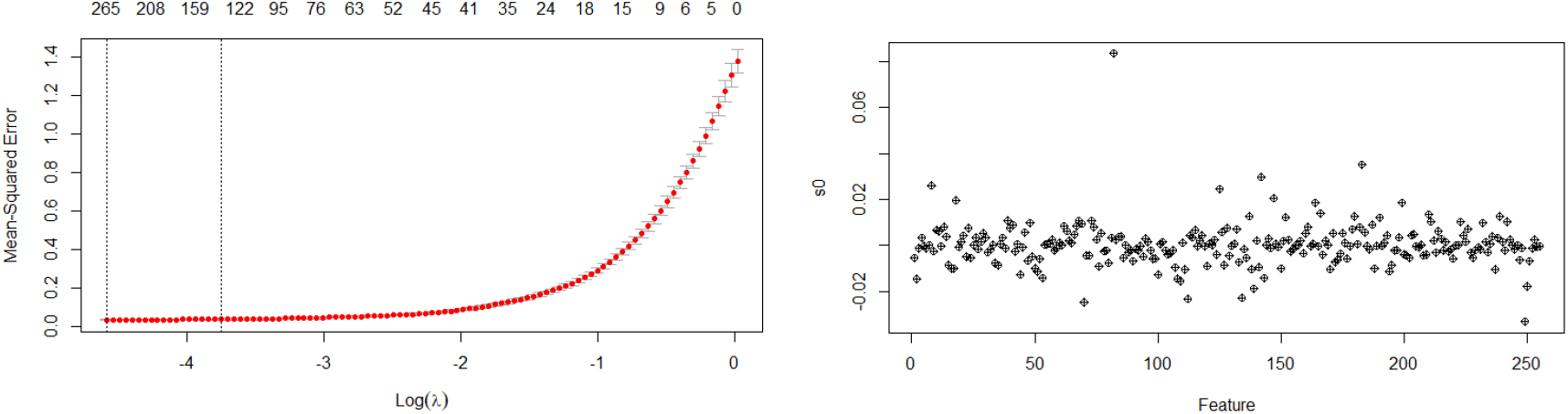
Calculates the mean squared error using K-fold cross-validation to suggest a lambda value for LASSO. Classification of the variables by mbLASSO regarding Lambda = 0.9 is 254.

Most features have small importance values (close to zero), meaning they do not significantly contribute to the model. A few features have higher values, suggesting they are the most important predictors. Sparse distribution: Many features might be eliminated (coefficients shrink to zero due to LASSO regularization).

In all these plots of Mean-Squared Error (MSE) versus Log(λ), likely from a cross-validation process in a mb-LASSO. The x-axis represents the logarithm of the regularization parameter (Log(λ)). The y-axis represents the Mean-Squared Error (MSE). The red points indicate MSE values for different values of λ. The error bars represent the variability (standard error) in MSE. The top axis values likely correspond to the number of nonzero coefficients in the model at each λ. The two vertical dotted lines might indicate the optimal λ values, commonly the minimum MSE (left) and the one-standard-error rule selection (right). λ_min (Left Line): Produces the most accurate model with minimal MSE. Retains more features (less regularization). λ_1se (Right Line): Provides a simpler model with fewer features but still within acceptable error limits. Often preferred for better generalization (reduces over fitting). The Classification for variables under lambda 0.2 is three hundred twenty one variables. These variables are effective for the RNA-Seq gene expressions. When we have lambda in minimum position then it will be shown that highest no of features are directly effects positively or negatively to the gene expression. This plot suggests that only a few variables contribute significantly, and many features could be discarded without losing much information. This is useful for feature selection, reducing dimensionality, and improving model efficiency. This scatter plot likely represents the importance scores of features after applying a variable selection method (e.g., LASSO, Ridge, or another feature selection technique). X-axis (Feature Index): Each point represents a different feature from the dataset, indexed numerically. Y-axis (s0 -Feature Importance Score or Coefficient Value): Indicates the importance or contribution of each feature to the model. Values close to zero suggest the feature is less relevant. Higher values indicate stronger influence in the model.

The feature selection in our study, so as the value of lamba increases the optimal lambda values gets decreasing as shown in Figure. Increasing the value of lambda intensifies the penalty applied to the model coefficients, causing a greater number of them to be pushed closer to zero. This process diminishes the influence of certain features or completely removes them from the model, thereby enabling automatic feature selection. In contrast, lowering lambda decreases the strength of the penalty, allowing more coefficients to remain substantial and preserving a larger set of features within the model.

Most features have small importance values (close to zero), meaning they do not significantly contribute to the model. A few features have higher values, suggesting they are the most important predictors. Sparse distribution: Many features might be eliminated (coefficients shrink to zero due to LASSO regularization).

In the above table there are the comparisons on lambda values ranges from 0.1 to 0.9. As the lambda increasing the optimal lambda turns down, whereas the selected variables make same scenario with optimal lamda. This table provides the relationship between tuning parameter (λ) and the number of selected features in a LASSO regression model. As λ increases, more regularization is applied, leading to fewer selected variables (feature elimination). At lower λ (0.1), more features are retained (386 selected). As λ increases, the number of selected features decreases, showing the effect of LASSO shrinking coefficients toward zero. At λ = 0.9, only 254 features remain, meaning LASSO has eliminated over 130 variables compared to λ = 0.1. There is a slight increase at λ = 0.7, which may indicate a model instability or numerical adjustment.

As λ increases, variables with smaller contributions to the model are more likely to have their coefficients shrunk to zero. Important variables (those with larger absolute coefficients) are retained longer as λ increases. Lambda helps prioritize variables by their relevance to the response as shown in Figure 15.

**Figure 15:**
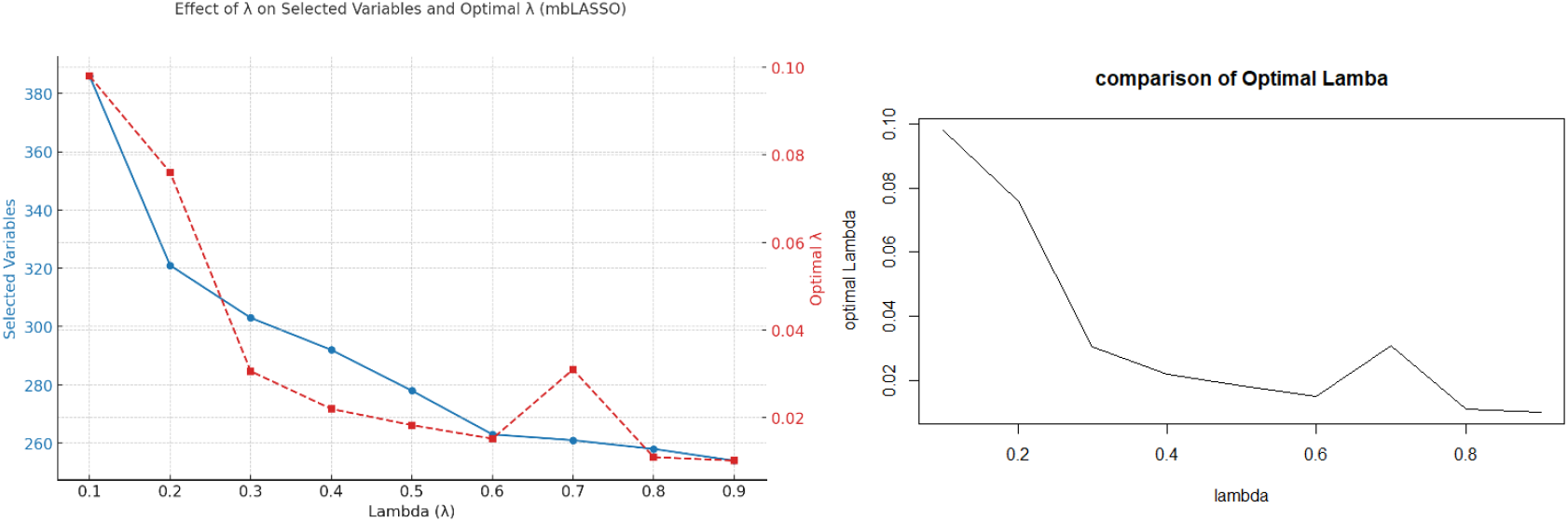
Comparison of Lambda values and the Optimal Lambda on different stages for λ=0.1-0.9

**Figure 16:**
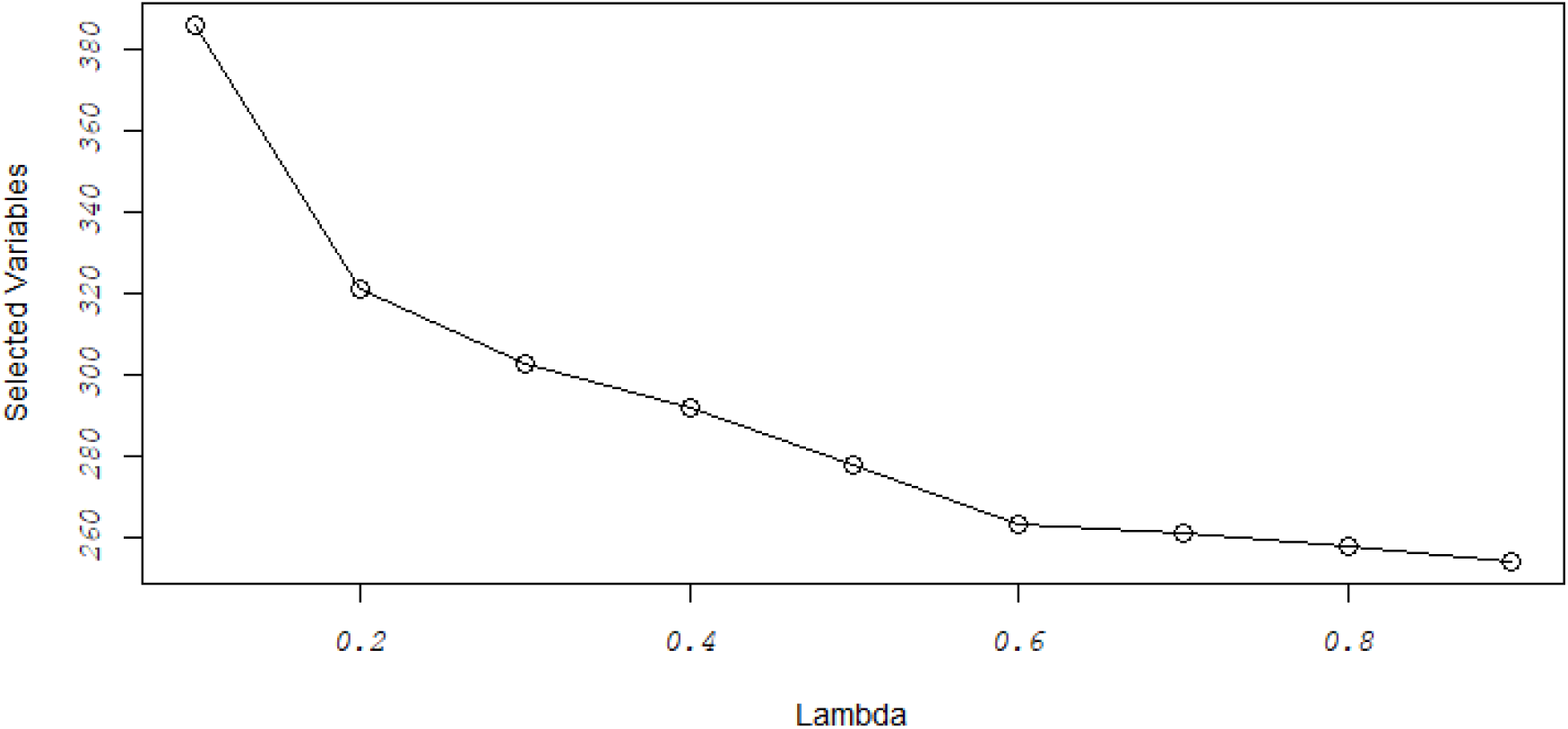
Feature Selection from multiple variables of RNA-Seq besides the lambda values 0.1-0.9. The number of variables decreases as λ is going in increasing trend

## 4. CONCLUSIONS

We have developed a generalized variable selection framework that focuses on selecting clusters of correlated variables by employing a multiblock Least Absolute Shrinkage and Selection Operator (mbLASSO) approach. This method effectively captures the structured relationships within grouped variables, enabling more meaningful and stable feature selection in high-dimensional data settings. The derived results could provide better biological understanding by selecting stable clusters where correlated variables are connected, which may lead to improved classification. As the lambda increasing the optimal lambda turns down, whereas the selected variables make same scenario with optimal lamda. This table provides the relationship between the tuning parameter (λ) and the number of selected features in mbLASSO model. As λ increases, more regularization is applied, leading to fewer selected variables (feature elimination). At lower λ (0.1), more features are retained (386 selected). As λ increases, the number of selected features decreases, showing the effect of LASSO shrinking coefficients toward zero. At λ = 0.9, only 254 features remain, meaning LASSO has eliminated over 130 variables compared to λ = 0.1. There is a slight increase at λ = 0.7, which may indicate a model instability or numerical adjustment. The feature selection in our study, so as the value of lamba increases the optimal lambda values gets decreasing. Higher lambda values strengthen the penalty on the model coefficients, driving more of them toward zero. This reduces the impact of some features or eliminates them entirely, effectively performing automatic feature selection. Conversely, lower lambda values lessen the penalty’s influence, allowing more features to stay in the model and contribute to its predictions.

